# Fast simulation of identity-by-descent segments

**DOI:** 10.1101/2024.12.13.628449

**Authors:** Seth D. Temple, Sharon R. Browning, Elizabeth A. Thompson

## Abstract

The worst-case runtime complexity to simulate haplotype segments identical by descent (IBD) is quadratic in sample size. We propose two main techniques to reduce the compute time, both of which are motivated by coalescent and recombination processes. We provide mathematical results that explain why our algorithm should outperform a naive implementation with high probability. In our experiments, we observe average compute times to simulate detectable IBD segments around a locus that scale approximately linearly in sample size and take a couple of seconds for sample sizes that are less than ten thousand diploid individuals. In contrast, we find that existing methods to simulate IBD segments take minutes to hours for sample sizes exceeding a few thousand diploid individuals. When using IBD segments to study recent positive selection around a locus, our efficient simulation algorithm makes feasible statistical inferences, e.g., parametric bootstrapping in analyses of large biobanks, that would be otherwise intractable.

## 1 Introduction

Simulation is a powerful tool in population genetics to forecast the genetic impact of evolutionary scenarios, perform statistical inference on models and their parameters, and develop and evaluate new methods (Hoban et al., 2012; Yuan et al., 2012). There are two main frameworks for population genetics simulations, each having its own use cases, advantages, and disadvantages. Forward simulation models the dynamics of entire populations over time regarding individuals and their interactions (Haller and Messer, 2019). This flexible approach can incorporate complex dynamics of selection, migration, and spatial context, among other features, at the cost of additional computation. Backward simulation models the genealogy of present-day samples strictly through their common ancestors and is less computationally intensive (Hoban et al., 2012).

The speed of backward simulation is in large part due to coalescent theory (Kingman, 1982a,b), which approximates the Wright-Fisher (WF) process (Wright, 1931) when the sample size is much smaller than the population size. The Kingman coalescent has been extended to address examples of migration (Nath and Griffiths, 1993), recombination (Hudson, 1983; Hudson and Kaplan, 1988), selection (Hudson and Kaplan, 1988; Kaplan et al., 1988), and demography (Hein et al., 2005). With recombination, the model becomes a sequence of correlated coalescent trees called the ancestral recombination graph (ARG). In recent years, numerous coalescent methods have been developed to simulate polymorphism data over large genomic regions efficiently (Ewing and Hermisson, 2010; Hudson, 2002; Kern and Schrider, 2016), having randomly placed mutations on tree branches at a fixed genome-wide rate. The `msprime` software is a popular and robust option for backward simulation that scales to entire chromosomes and thousands of individuals (Baumdicker et al., 2021). Hybrid frameworks with forward simulations (Haller et al., 2019) and standards set for species-specific simulations (Adrion et al., 2020; Lauterbur et al., 2023) have contributed to its widespread adoption.

Placing mutations on tree branches has linear complexity in sample size, which means analyses focusing on summary statistics of polymorphism data can be runtime inexpensive even in large samples. On the other hand, deriving the pairwise relationships between haplotypes is difficult for large sample sizes because the total number of computations scales quadratically in sample size. To be precise, two individuals share a haplotype segment identical-by-descent (IBD) if they inherit it from the same common ancestor. `msprime` has a feature to access IBD segments from the tree sequence, but its documentation warns that deriving and storing the IBD segments requires a lot of time and memory (Baumdicker et al., 2021). Another coalescent method `ARGON` simulates IBD segments as a feature within a much broader ARG-inference program (Palamara, 2016). These methods are the two current options to simulate IBD segments genome-wide in modestly sized samples. The runtime to simulate IBD segments with these programs has not been extensively benchmarked.

Long IBD segments can be informative about recent demographic changes (Browning and Browning, 2015; Browning et al., 2018; Cai et al., 2023; Palamara et al., 2012), recent positive selection (Browning and Browning, 2020; Temple et al., 2024), population-specific recombination rates (Zhou et al., 2020a), mutation rates (Tian et al., 2019), allelic conversions (Browning and Browning, 2024), rare variant association studies (Browning and Thompson, 2012; Chen et al., 2023), and close familial relatedness (Zhou et al., 2020c), whereas summary statistics like the fixation index *F*_*ST*_ (Weir and Cockerham, 1984) and Tajima’s *D* (Tajima, 1989) or models like the sequentially Markovian coalescent (SMC) (Li and Durbin, 2011), and its extensions (Schiffels and Durbin, 2014), concern population divergences and old selection events (Tajima, 1989; Weir and Cockerham, 1984), among other things. Methods using IBD segments thus serve as an important complementary approach to summary statistics and coalescent-based methods.

Distinguishing between alleles that are identical-by-state versus those that are identical-by-descent from a common ancestor can be challenging. Only those haplotypes extending over multiple centiMorgans, a unit of genetic distance to be defined in Section 2, can be detected as IBD with high accuracy (Freyman et al., 2021; Nait Saada et al., 2020; Naseri et al., 2019; Shemirani et al., 2021; Zhou et al., 2020b). We refer to IBD segments longer than a fixed Morgans threshold as “detectable”, where a user-defined threshold can depend on the dataset, the IBD segment detection method, and the tolerance to detection inaccuracies. Exceptionally long IBD segments are rare to observe outside of family studies, meaning that large sample sizes are required to observe enough for IBD-based analyses in outbred population studies.

Some methods require IBD data for the entire chromosomes (Browning and Browning, 2015; Palamara et al., 2012; Temple, 2024; Zhou et al., 2020a,c), which simulators like `msprime` (Baumdicker et al., 2021) and `ARGON` (Palamara, 2016) are suited for. Other statistical inferences concern estimator consistency (Temple et al., 2024), uncertainty quantification (Temple et al., 2024), and convergence to an asymptotic distribution (Temple and Thompson, 2024) around a single locus. Validating such theoretical results involves enormous simulations, for which `msprime` and `ARGON` are less suited.

In this work, we propose an algorithm to simulate IBD segments overlapping a focal location that is fast enough to validate asymptotic properties like consistency, confidence interval coverage, and weak convergence (Casella and Berger, 2002). We modify a naive approach (Temple et al., 2024), and then we argue that our modified approach should drastically decrease runtime with high probability. We demonstrate in some simulation examples that the modified algorithm’s average runtime scales approximately linearly with sample size, not quadratically.

## 2 Preliminary material

Backward simulation of IBD segment lengths overlapping a focal location involves two waiting time distributions: the time until a common ancestor and the genetic length until a crossover. Figure 1 illustrates the coalescent and recombination processes. Here, we formally define a parametric model for IBD segments overlapping a specific locus in terms of these processes.

**Fig 1.**
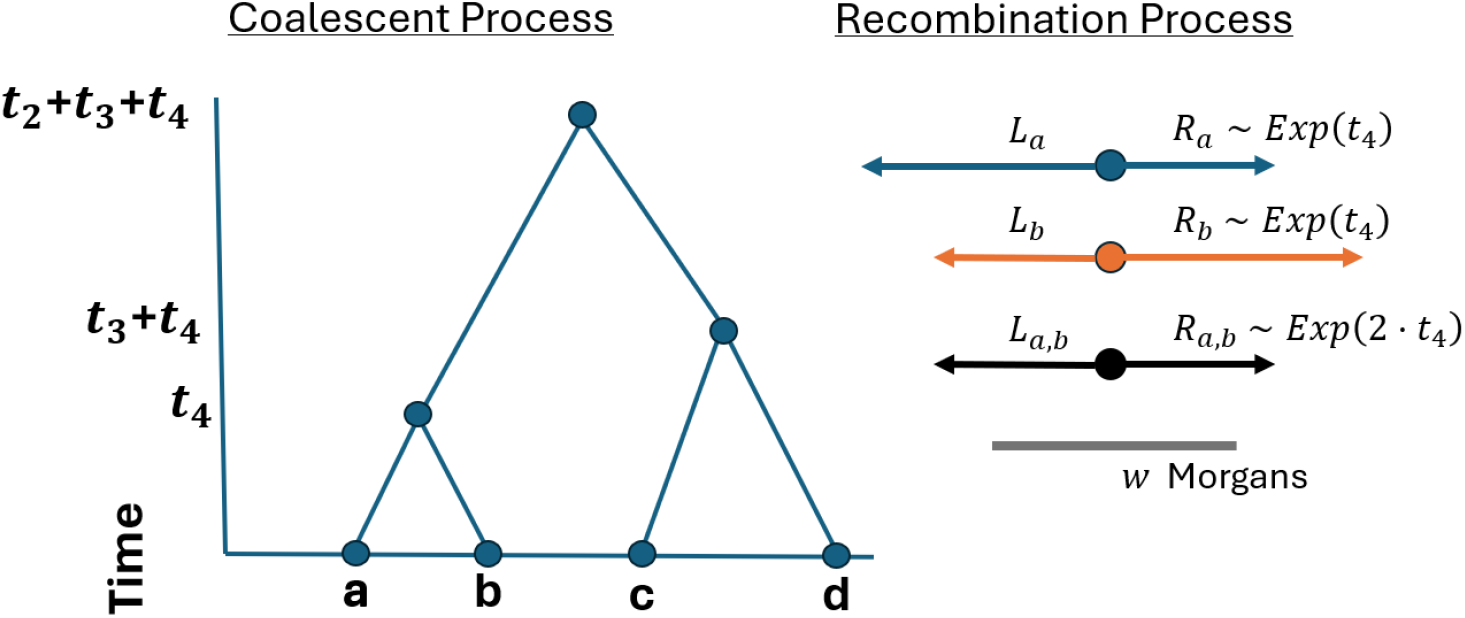
Conceptual framework for IBD segment lengths. (Left) Sample haplotypes *a, b, c, d* trace their lineages back to common ancestors at times *t*_4_, *t*_4_ +*t*_3_, *t*_4_ +*t*_3_ +*t*_2_. (Right) Relative to a focal point, the haplotype segments lengths *R*_*a*_, *R*_*b*_, *L*_*a*_, *L*_*b*_ are independent, identically distributed Exponential(*t*_4_). The lengths shared IBD are *R*_*a,b*_ := min(*R*_*a*_, *R*_*b*_) and *L*_*a,b*_ := min(*L*_*a*_, *L*_*b*_). The IBD segment length *W*_*a,b*_ := *L*_*a,b*_ + *R*_*a,b*_ ∼ Gamma(2, 2 · *t*_4_) exceeds the detection threshold *w* Morgans.

### 2.1 The time until a common ancestor

Let *n* be the haploid sample size, *k* ≤ *n* the size of a subsample, *N* (*t*) the population size *t* generations ago. Unless otherwise specified, time *t* ≥ 0 always refers to time backward from the present day. For constant population size, note that *N* = *N* (*t*) for all *t*. In the discrete-time Wright-Fisher (WF) process, each haploid has a haploid ancestor in the previous generation. If haploids have the same haploid ancestor, their lineages join.

Let the random variable *T*_*k*_ denote the time until a common ancestor is reached for any two of *k* haploids. The random variable 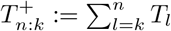 is the time until *n* − *k* + 1 coalescent events. The time to the most recent common ancestor (TMRCA) of the sample is 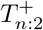. The probability that the time until the most recent common ancestor of two specific haploids is

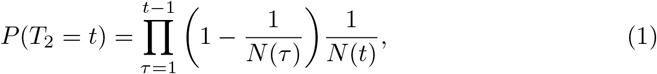

where 1*/N* (*τ*) is the probability that a haploid has the same haploid parent as the other haploid at generation *τ*. The approximate probability that the time until a common ancestor is reached for any two of *k* haploids is

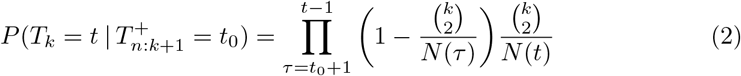

when *k* is much smaller than min_*t*_ *N* (*t*) (Hein et al., 2005). The geometric model assumes that multiple coalescent events in a single generation are improbable. Its rate 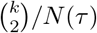 is the probability that any two of *k* haploids have the same haploid parent at generation *τ*.

The Kingman coalescent (Kingman, 1982b,a) comes from the continuous time limit of Equations 1 and 2 for large constant population size *N*. Specifically, *T*_*k*_ converges weakly to Exponential (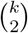) for *k* ≪ *N, N* → ∞, and time is scaled in units of *N* generations. Henceforth, we consider the positive real-valued *T*_*k*_ in units of *N* generations. Varying population sizes *N* (*t*) are implemented by rescaling time *post-hoc* in a coalescent with constant population size *N* (Hein et al., 2005).

### 2.2 The distance until crossover recombination

The genetic distance between two points is the expected number of crossovers between them in an offspring gamete. This unit of haplotype segment length is the Morgan. Assuming no interference in double-stranded breaks and that crossovers occur randomly and independently, Haldane (1919) derives that the genetic distance until crossover recombination is exponentially distributed, with the Poisson process modeling the crossover points along the genome. The number of crossovers between two points is then Poisson distributed with mean equal to the genetic distance between the two points, which leads to the Haldane map function connecting Morgans to the recombination frequency. (The Haldane map function is *ρ* = 0.5(1 − exp(−2*d*)), where *ρ* is the recombination frequency and *d* is the genetic distance.)

From a fixed location, the Morgan distance until a crossover in one gamete offspring is distributed as Exponential(1). An important property of the exponential random variable is that the minimum of independent exponential random variables is an exponential random variable with a rate that is the sum of the rates of the independent random variables. Since meioses are independent after *t* meioses the haplotype segment length to the right of a focal location is distributed as Exponential(*t*), where *t* is the rate parameter.

Let *a* and *b* be sample haplotypes in the current generation. Define *L*_*a*_, *R*_*a*_ | *t* ∼Exponential(*t*) to be sample haplotypes *a*’s recombination endpoints to the left and right of a focal location. Since crossovers to the left and right are independent, the extant width derived from the ancestor at time *t* is *W*_*a*_ := *L*_*a*_ + *R*_*a*_ | *t* ∼ Gamma(2, *t*).

Because recombination events are independent in the *t* meioses descending to *a* and *b* from their common ancestor, the IBD segments that are shared by *a* and *b* are *L*_*a,b*_, *R*_*a,b*_ | *t* ∼ Exponential(2*t*) and *W*_*a,b*_ | *t* ∼ Gamma(2, 2*t*). Under this model, the lengths of IBD segments are thus shorter, with a higher probability the more removed its common ancestor is from the present day. This fact is a key motivation for the fast algorithm we develop.

## 3 An efficient algorithm to simulate identity-by-descent segments

Based on Sections 2.1 and 2.2, the blueprint to simulate IBD segment lengths around a locus is as follows: 1) simulate a coalescent tree for a sample from a population, 2) draw recombination endpoints to the left and right of a focal point at each coalescent event, and 3) derive from the recombination endpoints the haplotype segment lengths that are shared IBD. The third step involves calculating the minimum lengths to the right and left of a focal point for every pair of haplotypes, which is the computational bottleneck in simulating IBD segment lengths. Making fewer haplotype comparisons, without sacrificing the exactness of simulation, is the way to decrease compute times. In Algorithm 1, we state the method to simulate long IBD segments around a single locus. We make four modifications to the naive simulation algorithm, which are designed to reduce compute times when the primary goal is to generate IBD segments longer than some detection threshold. These implementations reduce compute times due to the mathematical properties of the coalescent time and recombination endpoint distributions.

First, whenever there is likely to be more than one coalescent event in a Wright-Fisher (WF) generation, we approximate the sampling of haploid parents as a binomial random variable (Section 4). Second, we exchange the Kingman coalescent for the discrete-time WF model once the number of non-coalesced haploids is much smaller than the population sizes. This implementation is similar to the hybrid simulation approach in Bhaskar et al. (2014). Third, we do not consider a sample haplotype for IBD segment calculation at future coalescent events once its haplotype segment length is less than the specified detection threshold, which we refer to as “pruning”. In Section 5, we elaborate on the rare probability of long haplotype segments in large populations. Fourth, we combine two sample haplotypes for IBD segment calculation at future coalescent events if they share the same left and right recombination endpoints, which we refer to as “merging”. In Section 6, we derive results concerning the probability of merging. We implement pruning and merging using object-oriented programming.

## 4 An approximation of the Wright-Fisher process in large samples

Simulating the Kingman coalescent is much faster than simulating the discrete-time WF process. The accuracy of the Kingman coalescent requires that the sample size is much smaller than the population size. This requirement is so that the probability of there being more than one coalescent event in a generation is small. The assumption that the sample size is small relative to the population size can be violated in analyses of human biobanks. Under this violation, the coalescent approximation can deviate significantly from the exact discrete-time WF model (Bhaskar et al., 2014; Palamara, 2016; Wakeley and Takahashi, 2003).

In the following approximations for the sampling of haploid parents at each generation, we suppress the dependence on the generation time *t*. Let *k* := *k*(*t* − 1) be the number of lineages at generation *t* − 1. Let *k*^*′*^ := *k*(*t*) and *N* ^*′*^ := *N* (*t*) be the number of lineages and the population size in the previous generation *t*. The probability that a parent among {1, …, *N* ^*′*^} has no children is (1 − 1*/N* ^*′*^)^*k*^. The probability that a parent has at least one child is 1 − (1 − 1*/N* ^*′*^)^*k*^. The Taylor series expansion in 1*/N* ^*′*^ about zero is

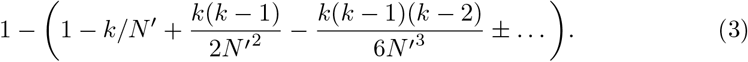

### Algorithm 1

Efficient simulation of IBD segment lengths

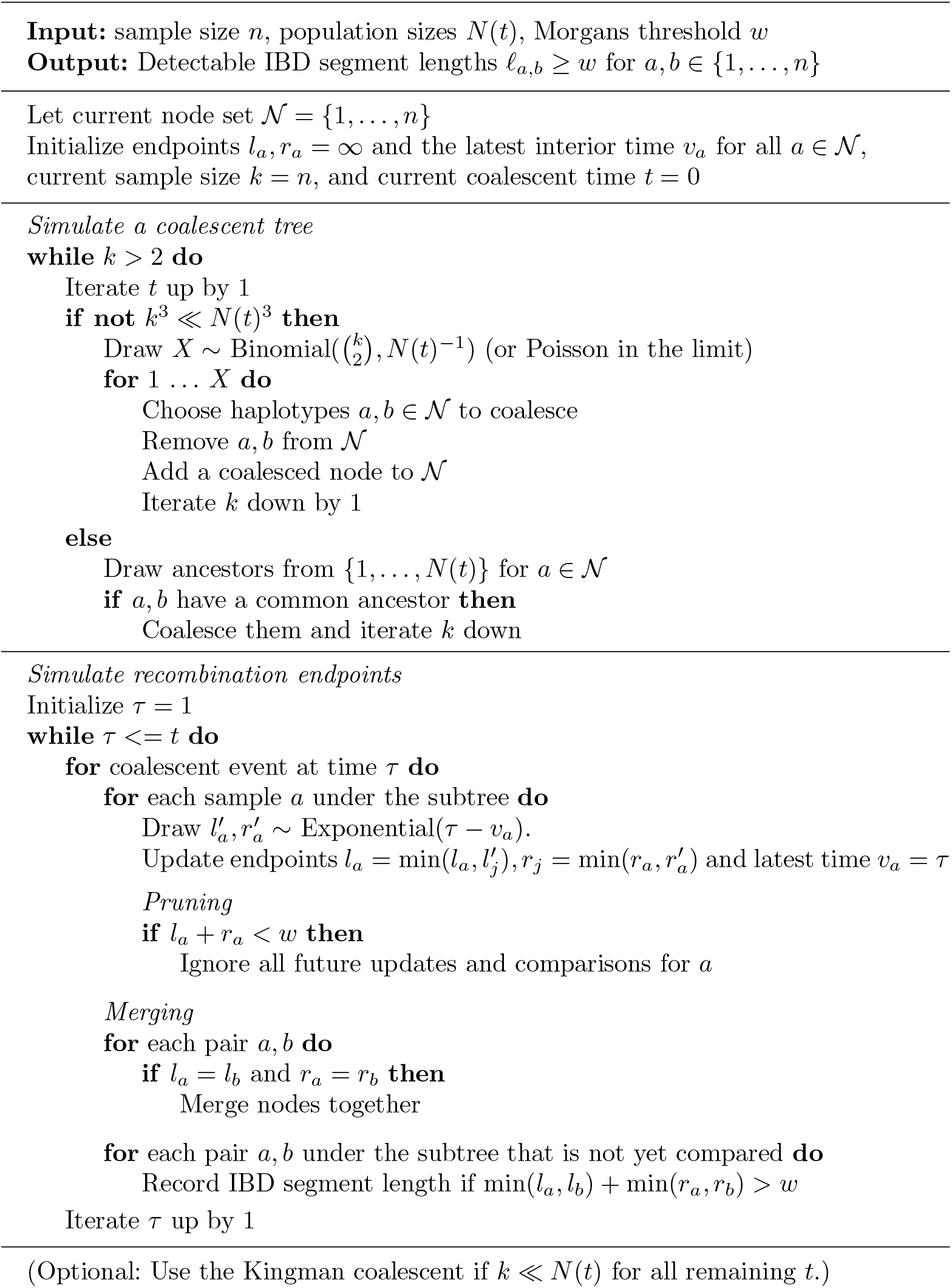

The second order approximation 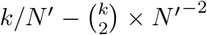 is accurate if *k*^3^ = *o*(*N* ^*′*3^). The expected number of parents in the previous generation *t* with a child in generation *t* − 1 is then

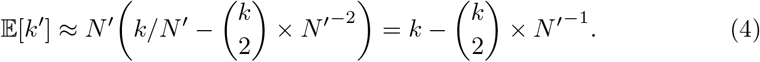

As an example, consider a sample of twenty thousand haploids whose ancestral population sizes in the recent ten generations are more than two hundred thousand haploids. The second order approximation is accurate for the first ten generations because *k*^3^ · *N* ^−3^ = 10^−3^ when the sample size *k* = 2 · 10^4^ is an order of magnitude smaller than the population size *N* = 2 · 10^5^. For this choice of *k* and *N* ^*′*^, the expected number of coalescent events per generation is approximately five hundred.

Compared to drawing a parent for each child and then scanning a vector of size *k* for siblings, simulating the number of coalescent events in one generation from Binomial 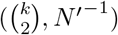 can be an efficient approximation. The last term being subtracted in Equation 4 is equal to the expected value of a Binomial random variable of 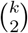 trials with success probability *N* ^*′*−1^. Next, let *A*_1_ and *A*_2_ be the number of children from two specific haploid parents among the *N* ^*′*^ parents in the previous generation. If *A*_1_ and *A*_2_ are independent, then *P* (*A*_1_ = *a*_1_, *A*_2_ = *a*_2_) = *P* (*A*_1_ = *a*_1_) × *P* (*A*_2_ = *a*_2_). *A*_1_ and *A*_2_ are not independent, but the difference between the left term *P* (*A*_1_ = *a*_1_, *A*_2_ = *a*_2_) and the right term *P* (*A*_1_ = *a*_1_) × *P* (*A*_2_ = *a*_2_) can be vanishingly small when *N* ^*′*^ is large. The probability *P* (*A*_1_ = *a*_1_, *A*_2_ = *a*_2_) is derived by choosing *a*_1_ among *k* samples to have the same parent and then choosing *a*_2_ among *k* − *a*_1_ samples to have a same parent distinct from the parent of the first *a*_1_ samples.

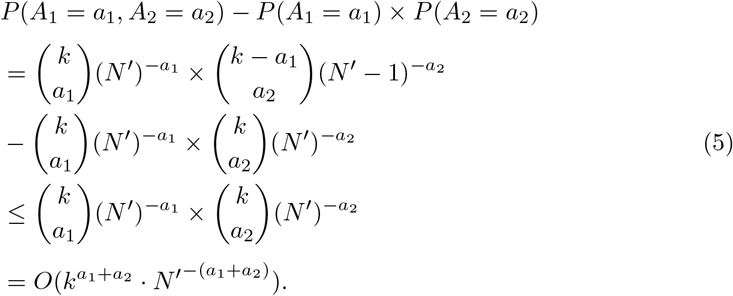

If both *A*_1_ and *A*_2_ have two or more children (min(*a*_1_, *a*_2_) ≥ 2), then Equation 5 is *o*(1) when the second order approximation *k*^3^ = *o*(*N* ^*′*3^) is accurate.

In Algorithm 1, we assume that all simultaneous coalescent events are the result of only two children having the same parent. Bhaskar et al. (2014) have shown that the majority of simultaneous coalescent events in a generation are of this type. Due to the coalescent and WF approximations, our method is not exact with respect to the time until a common ancestor.

## 5 The probability of detectable haplotype segment lengths

Within tens of generations, most haplotype segment lengths are shrunk by crossovers to a genetic length less than detection thresholds that are used in IBD-based analyses. A Morgans length threshold at least greater than 0.01 is typical in applied research (Browning and Browning, 2015, 2020; Temple et al., 2024; Tian et al., 2019; Zhou et al., 2020a). The probabilities of a detectable haplotype segment to the right of and overlapping a focal location, *R*_*a*_ and *W*_*a*_, respectively, conditional on coalescent time *Nt* (in generations), are

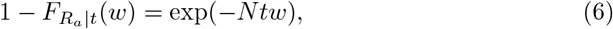

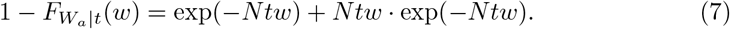

Figure S1 shows that the upper tail probabilities of *R*_*a*_ and *W*_*a*_ are decreasing exponentially over *Nt* generations. The probabilities of haplotype segment lengths greater than 0.01 can be far from zero when the haplotype is descendant from an ancestor within the last 100 generations. The probabilities of haplotype segments lengths greater than 0.02 are nearly zero when they are descendant from an ancestor more than 300 generations ago. (But exponential random variables have heavy upper tail probabilities, so, in large samples, we may detect some long IBD segments descendant from ancestors older than 300 generations.)

For large populations, the coalescent times of ancestral lineages can be much greater than 500 generations. The expected time of the (*n* − *k* + 1)^th^ coalescent event can be derived as:

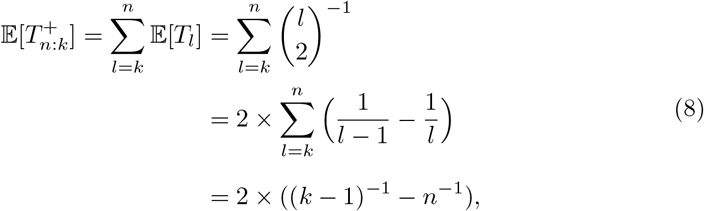

where 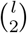 is the rate parameter for the time until a common ancestor is reached for any two of *l* haploids. For *N* = 10, 000 and *n* → ∞, the expected coalescent time 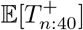 is 512.82 generations. For *N* = 100, 000 and *n* → ∞, the expected coalescent time E[*T*_*n*:400_] is 501.25. If many recombination endpoint comparisons happen at the coalescence of common ancestors older than five hundred generations ago, many haplotypes can be pruned ahead of time. The pruning technique does not compromise the exactness of simulating IBD segment detectable beyond a length threshold.

## 6 The probability that recombination endpoints are shared between haplotypes

At some point in the past, two sample haplotypes may share the same recombination endpoints to the left and right of a fixed location. Without loss of generality, let haplotypes *a* and *b* coalesce to their common ancestor *c* at time *u*, and let haplotypes *c* and *d* coalesce to their common ancestor *e* at time *u* + *v*. Figures S2 and S3 illustrate the coalescent tree in this scenario. Observe that the recombination endpoints to the right *R*_*a,c*_, *R*_*b,c*_ ∼ Exponential(*u*) and *R*_*c,e*_ ∼ Exponential(*v*).

The merging step in Algorithm 1 serves to avoid comparing both the endpoints of *a* and *b* with *d* when *a* and *b* have the same endpoints at time *u* + *v*. Specifically, if *a* and *b*’s shared recombination endpoint *R*_*c,e*_ is smaller than their separate endpoints *R*_*a,c*_ and *R*_*b,c*_, we can henceforth treat them as the same haplotype without loss of information (Figure S2). If either of the individual lengths *R*_*a,c*_ or *R*_*b,c*_ are smaller than the common length *R*_*c,e*_, we cannot merge the haplotypes without losing information (Figure S3).

The probability that haplotypes *a* and *b* have the same recombination endpoint at time *u* + *v* is *v*(2*u* + *v*)^−1^. We derive a result that replaces arbitrary coalescent times *u* and *u* + *v* with double the expected times after the (*n* − *k*)^th^ and (*n* − *j*)^th^ coalescent events, respectively.

### Proposition 1.

*Let* 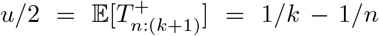 *and* 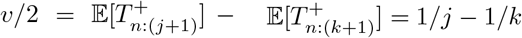 *(Equation 8). For j* = *o*(*k*),

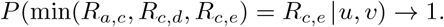

*Proof*. Note that *j* = *o*(*n*) as well because *k* ≤ *n*.

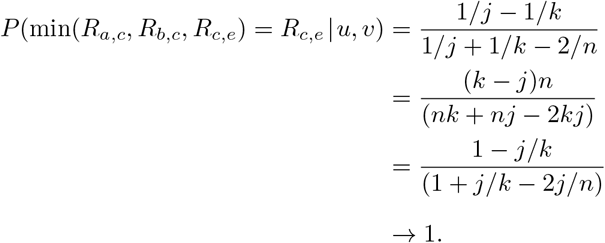

The implication of Proposition 1 is that haplotypes that share a recent common ancestor should have the same endpoints at the most distant common ancestors.

Since recombinations to the right and left of a focal location are independent, the result of Proposition 1 extends to simulating IBD segments overlapping a focal location. Figures S2 and S3 illustrate that merging occurs when the minimum recombination endpoints to the left and right of the focal location are drawn for the common ancestor *c* (min(*R*_*a,c*_, *R*_*b,c*_, *R*_*c,e*_) = *R*_*c,e*_ and min(*L*_*a,c*_, *L*_*b,c*_, *L*_*c,e*_) = *L*_*c,e*_).

## 7 The number of identity-by-descent comparisons

Pruning and merging should be most effective at reducing runtime if the majority of recombination endpoint comparisons happen at the oldest coalescent events. For these oldest coalescent events, we show that without pruning nor merging the expected number of IBD comparisons is of the same order as the worst-case number of IBD comparisons, which is asymptotically equivalent to the sample size squared.

Consider a random bifurcating tree. Here, and nowhere else, we work downward from the root of the tree. Throughout, we assume that *n* equals a power of 2 to simplify the floor and ceiling functions ⌊*n/*2^*j*^⌋ = ⌈*n/*2^*j*^⌉ for *j* ∈ N. At the coalescent event *T*_2_, the tree bifurcates into two subtrees. At the coalescent event *T*_3_, the scenario with the worst case number of comparisons is subtrees of size *n/*2, *n/*4, and *n/*4. In general, at each coalescent event, the worst case is to split in half the largest subtree, depicted in Figure S4A.

Let *B*_*j*_ be the size of one subtree randomly bifurcated from a subtree of size *B*_*j*−1_. Figure S4 illustrates these subtree sizes in the context of a random bifurcating tree. The number of recombination endpoint comparisons is *B*_*j*_(*B*_*j*−1_ − *B*_*j*_). In Theorem 2, we relate the expected value and covariance of *B*_*j*_(*B*_*j*−1_ − *B*_*j*_) to the worst-case *n/*2^2*j*^ computations. The result concerns a bounded number of standard deviations from the expected value, which is a stronger notion than the expected number of computations Θ(·). The intuition is that a Binomial(*m*, 1/2) random variable’s coefficient of variation *m*^−1*/*2^ converges to 0 as *m* gets large. The general proof strategy is to recursively apply the law of total covariance and identify the exponents in the dominating terms.

### Theorem 2.

*Let B*_*j*_ ∼ *Binomial*(*B*_*j*−1_, 1*/*2) *for bounded index j* ≥ 1 *and B*_0_ = *n*.

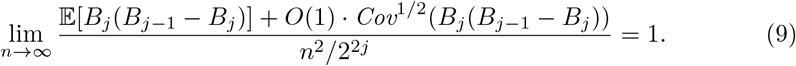

*Proof*. We must calculate the expected value and the covariance in the numerator. Let *B* ∼ Binomial(*m*, 1/2).

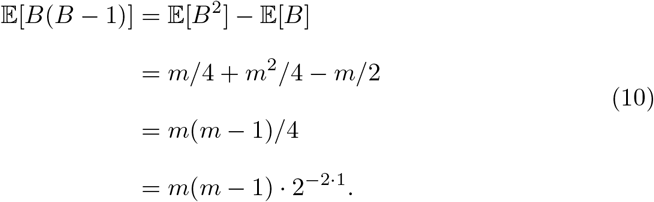

Using the law of total expectation, we solve the expected value for *j* = 2.

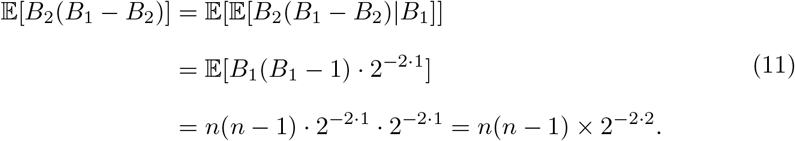

Applying Equation 10 recursively, we derive the general formula

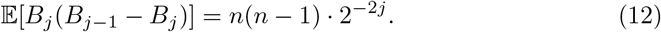

The limit of Equation 12 divided by *n*^2^ ·2^−2*j*^ is one. Next, we require that the standard deviation is of order less than *n*^2^. Using the law of total covariance, we derive in Lemma 3 that Cov(*B*_*j*_(*B*_*j*−1_ − *B*_*j*_)) ∼ *n*^3^, where ∼ means asymptotically equivalent. Consequently,

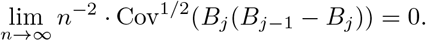

We remark that our marginal calculations along one branching path are not the same as deriving the expected number of comparisons at the final *j* coalescent events, the latter of which depends on the tree topology. Harding (1971) discusses the intractability of calculating probability masses for a tree topology with many leaves, which is a limiting factor in deriving the expected number of comparisons at the final *j* coalescent events. In Appendix A, we give moment calculations from Dahmer and Kersting (2015) that offer a complementary perspective on the number of IBD comparisons, reiterating that a number of computations ∼ *n*^2^ should occur at and near the root of the coalescent tree.

## 8 Empirical results

Temple et al. (2024) and Temple and Thompson (2024) use our algorithm to conduct enormous simulation studies involving sample sizes as large as ten thousand individuals and tens of millions of runs. (Individuals are “diploids”, which we implement as a haploid model with the number of haploids equal to the number of individuals times 2.) Their empirical studies are feasible because of the pruning and merging techniques, whose effects on runtime we benchmark in this section. We also benchmark runtimes for `msprime` and `ARGON`, showing that these existing methods to simulate IBD can take more than an hour to complete one run when sample size exceeds five thousand diploid individuals.

### 8.1 Experimental setup

#### 8.1.1 Demographic scenarios

We consider two complex demographic scenarios and constant population sizes. Figure S5 shows the demographic scenarios graphically. These demographic scenarios are the same as those used in Temple et al. (2024), Temple and Thompson (2024), and Temple (2024). We refer to the complex demographic scenarios as examples of three phases of exponential growth and a population bottleneck. The three phases of exponential growth scenario involves an ancestral population of five thousand individuals that grew exponentially at different rates in three different time periods. This demographic model is similar to the “UK-like” model in Cai et al. (2023). The population bottleneck scenario involves an ancestral population of ten thousand individuals that grew exponentially at a fixed rate but experienced an instantaneous reduction in size twenty generations before the present day.

#### 8.1.2 Hard selective sweeps

We also consider a genetic model for positive selection (Fisher, 1923; Haldane, 1924, 1932) that is described in Crow and Kimura (1970), Temple et al. (2024), and Temple (2024) as well as in many other articles. Briefly, the allele frequency *p*_*s*_(*t*) decreases backward in time as a function of a nonnegative selection coefficient *s*. The selection coefficient reflects the advantage the allele has relative to alternative alleles. The larger the selection coefficient is, the faster the allele frequency increased. Also, the larger the selection coefficient is, the more detectable IBD segments there are on average.

Positive selection around a locus is implemented via a coalescent with two subpopulations: one subpopulation has the sweeping allele, and one subpopulation does not have the sweeping allele. The population sizes are *N*_*e*_(*t*)·*p*(*t*) and *N*_*e*_(*t*)·(1−*p*(*t*)). Until the coalescent reaches the sweeping allele’s time of *de novo* mutation, IBD segments are not possible between individuals in separate subpopulations.

### 8.1 Compute times

#### 8.2.1 Simulating identity-by-descent segment lengths around a locus

To assess the effect of the pruning and merging rules, we evaluate four implementation strategies: merging and pruning (Algorithm 1), pruning only, merging only, and neither pruning nor merging (the naive approach). For each implementation, we run five simulations for sample sizes increasing by a factor of 2, recording the average wall clock compute time. The upper bound on sample size that we consider is 128,000 individiuals, which is of the same order as the UK Biobank data (Bycroft et al., 2018). Figure 2 shows the average runtime per sample size between the implementations.

**Fig 2.**
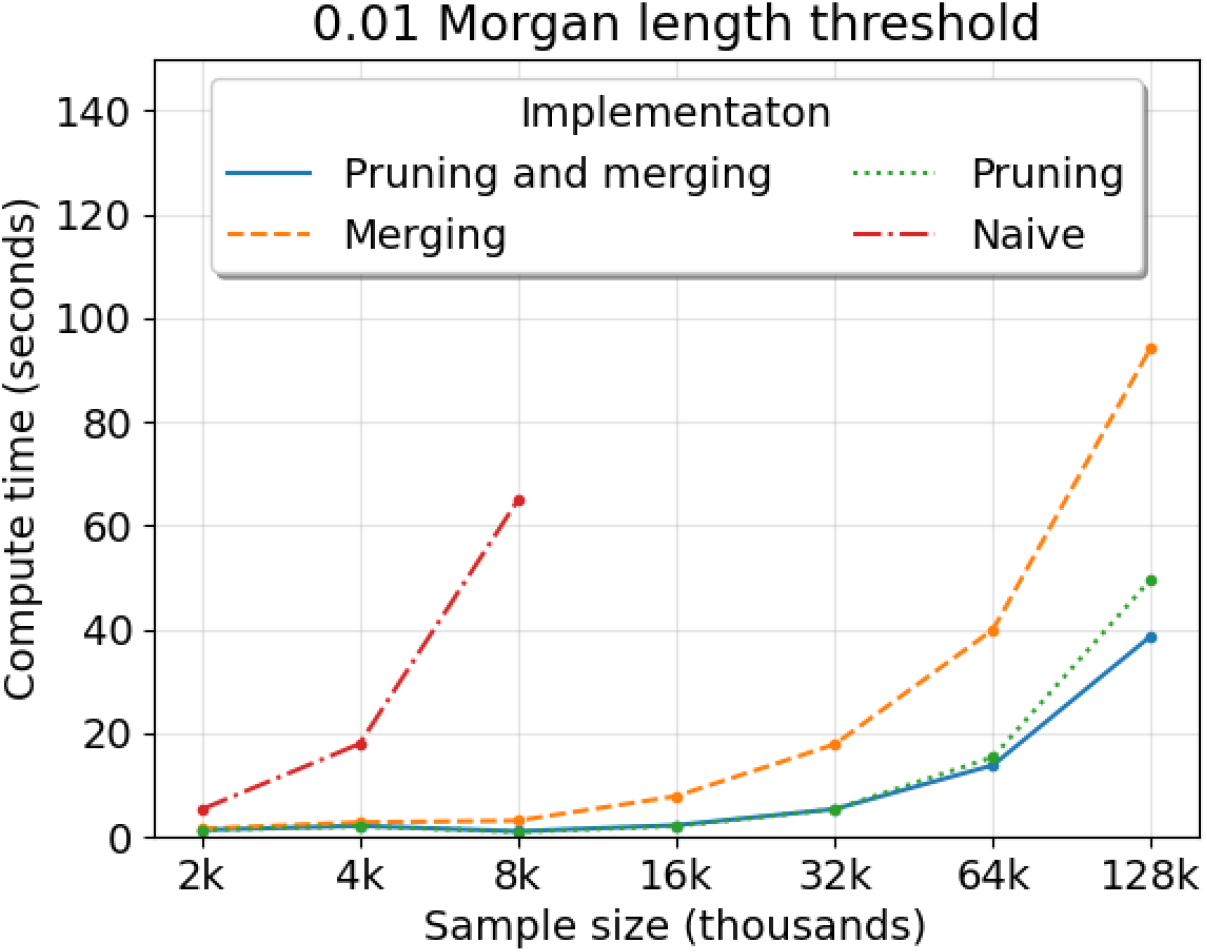
Compute time to simulate IBD segment lengths around a locus depending on algorithm implementation. Compute time (*y*-axis) in seconds by sample size (*x*-axis) in thousands is averaged over five simulations. The legend denotes colored line styles for implementations using Algorithm 1 as is (blue), merging only (orange), pruning only (green), and neither pruning nor merging (red). The main text describes “merging” and “pruning” techniques. The demography is the population bottleneck. The Morgans length threshold is 0.01.

Simulating IBD segment lengths without pruning nor merging takes more than one minute on average for eight thousand samples. Simulating IBD segment lengths with either pruning or merging can take less than one minute for sixty-four samples. Pruning appears to give a larger reduction in compute time than merging. Merging can further reduce runtime for sample sizes greater than one hundred thousand. The difference in five to ten seconds can be important when the number of simulations is enormous, as is the case in the Temple et al. (2024) and Temple and Thompson (2024) studies.

One important influence on runtime is the detection threshold. Figure 3A shows the algorithm’s average runtime per sample size for different detection thresholds on segment length. With the 0.0025 Morgans cutoff, the quadratic behavior of runtime is visually apparent when more than twenty thousand samples are simulated, whereas the trend is less obvious for detection thresholds greater than 0.0050 Morgans. The algorithm is at least twice as fast on average for detection thresholds ≥ 0.02 Morgans versus those ≤ 0.0050 Morgans.

**Fig 3.**
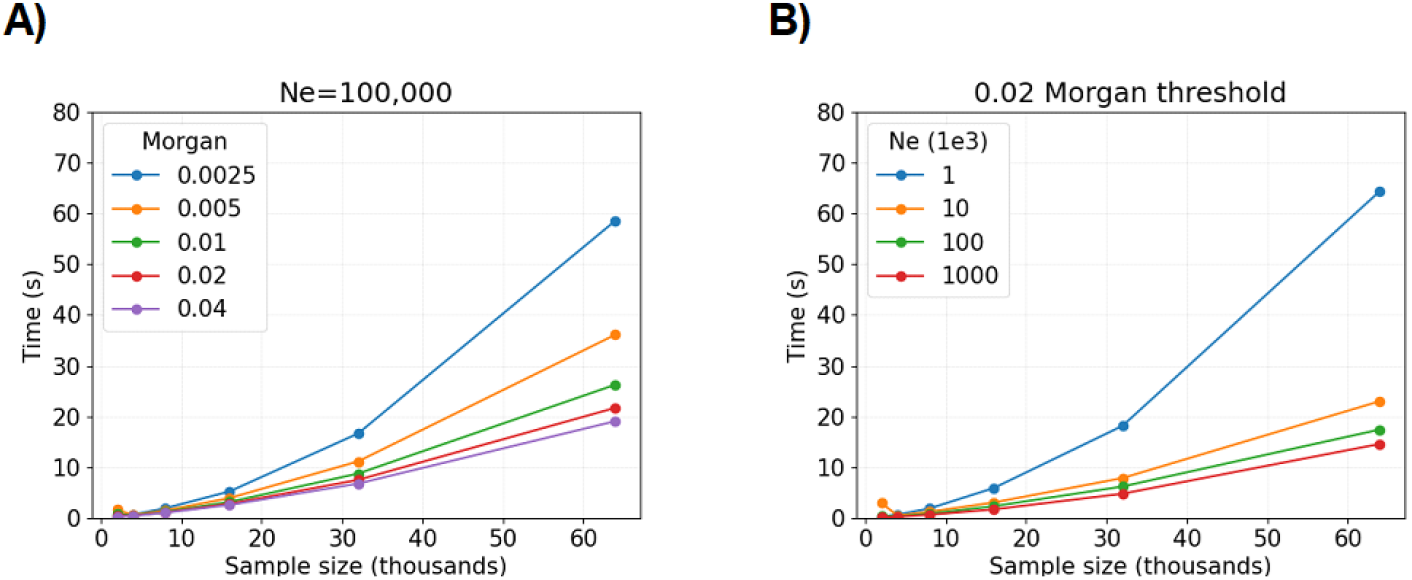
Compute time to simulate IBD segment lengths around a locus depending on the detection threshold and population size. Compute time (*y*-axis) in seconds by sample size (*x*-axis) in thousands is averaged over five simulations. The legends denote colored line styles for A) different detection thresholds (in Morgans) with *N* = 10^5^ fixed or B) different population sizes with 0.02 Morgans fixed.

Another important influence on runtime is the population size. Figure 3B shows the algorithm’s average runtime per sample size for different constant population sizes.

The algorithm is at least twice as fast on average for population sizes *N* ≥ 10, 000 versus *N* ≤ 1, 000. Population sizes are estimated to be at least ten thousand for many model organisms (Adrion et al., 2020; Lauterbur et al., 2023).

Figure S6 shows the algorithm’s average runtime per sample size for different demographic scenarios and varying selection coefficients. Simulating IBD segment lengths takes more time for the population bottleneck and three phases of exponential growth scenarios compared to constant-size population scenarios. Runtime increases with the selection coefficient. The highest average measurement is more than four minutes for sixty-four thousand samples, the population bottleneck scenario, and *s* = 0.04.

Now, we perform twenty simulations each for sample sizes 2 · 10^4^, 4 · 10^4^, 8 · 10^4^, 16 · 10^4^, and 32 · 10^4^ and regress on runtime. The linear models in runtimes **Y** ∈ ℝ, sample sizes **X** ∈ ℝ, and regression coefficients ***β*** be:

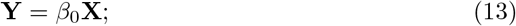

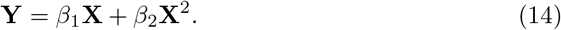

We measure the proportion of a fitted value explained by the linear effect as

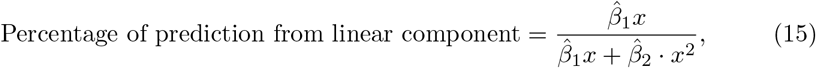

where *x* is a sample size. Figure 4 shows that the linear component in Equation 14 explains more than fifty percent of predictions for sample size ≤ 16 · 10^4^, which we say demonstrates approximately linear computational complexity in this domain. Next, we estimate 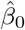 equal to 0.0913 and 0.0707 in Equation 13 for population sizes *N* = 10, 000 and *N* = 100, 000. We interpret this to mean that the expected runtime increases by 0.0913 and 0.0707 seconds for each one unit increase to sample size (in thousands), at least up to *n* ≤ 16 · 10^4^.

**Fig 4.**
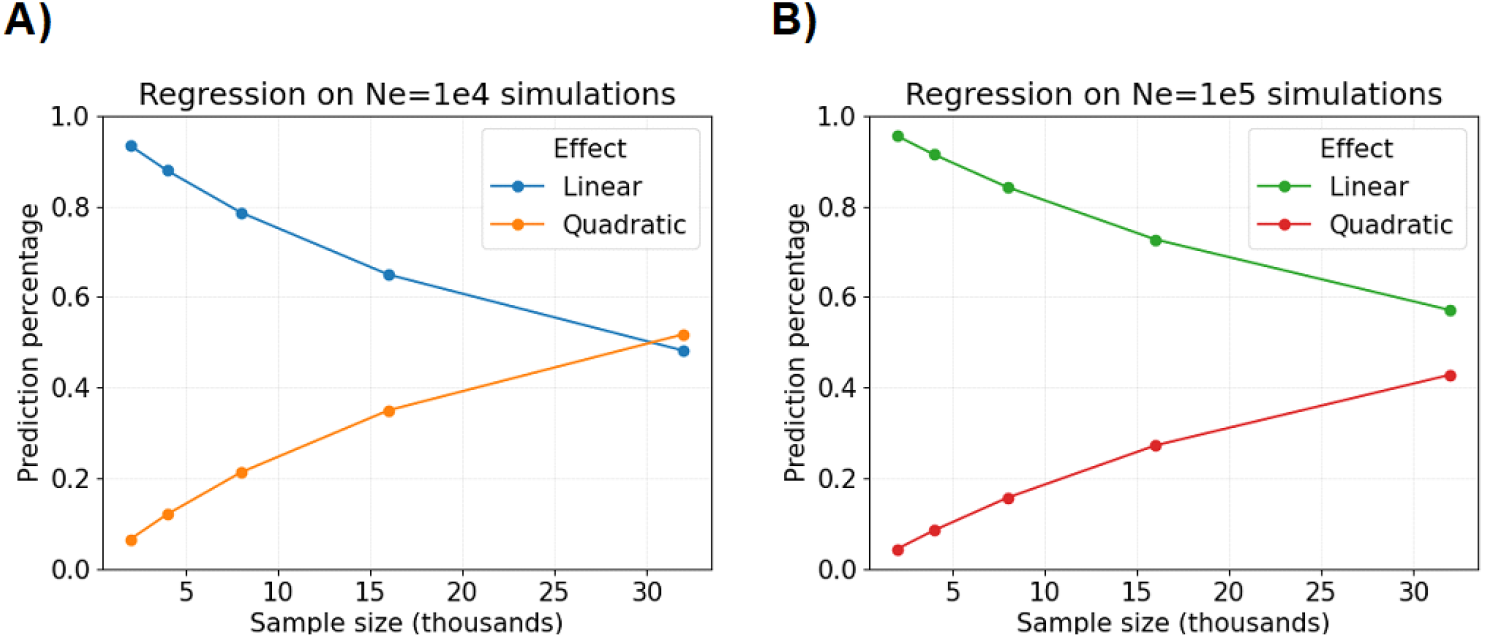
Percentage of regression model predictions explained by linear and quadratic effects. The percentage of predicted compute time (*y*-axis) in seconds by sample size (*x*-axis) in thousands with respect to linear and quadratic effects. Plots show results for A) the constant population size *N* ≡ *N*_*e*_ = 10, 000 versus B) *N*≡ *N*_*e*_ = 100, 000. The detectable IBD segments are simulated with a 0.02 Morgans threshold.

Overall, we benchmark that our simulation algorithm can be run tens of thousands of times within a day on one core processing unit of an Intel 2.2 GHz compute node. Despite performance savings, we observe that our simulation algorithm maintains quadratic behavior in sample size (Figure 2 and Figure S6). One explanation for this finding is that a sizeable fraction of all lineages coalesce at least once in the first few generations when the sample size exceeds ten thousand (Bhaskar et al., 2014).

#### 8.2.2 Simulating identity-by-descent segment lengths from the ancestral recombination graph

We measure the times it takes `ARGON` (Palamara, 2016) and `msprime` (Baumdicker et al., 2021) with `tskibd` (Guo et al., 2024) to simulate detectable IBD segments around a locus. The `tskibd` program concatenates short IBD segments from `msprime` tree sequences into detectable IBD segment lengths. To measure the computing time of these approaches, we do not include the time to simulate an ARG.

We simulate IBD segments ≥ 0.02 Morgans in a 0.07 Morgans region, which is a large enough region to contain all IBD segments ≥ 0.02 Morgans around its central location. Both programs visit nodes in the ARG in small, non-overlapping sliding windows. We consider window sizes of 0.0001 and 0.00001 Morgans in benchmarking runtimes. We compare compute times to those of Algorithm 1 with the 0.02 Morgans detection threshold.

Table 1 reports the average runtimes of each method for increasing sample size in the population bottleneck demographic scenario. `ARGON` takes nearly an hour to simulate the detectable IBD segment lengths of two thousand diploids. We do not run it for more than two thousand diploids due to concerns surrounding quadratic runtimes. With 0.0001 Morgans windows, `tskibd` takes less than twenty minutes to simulate the detectable IBD segment lengths of four thousand diploids and a little over an hour to simulate the IBD length distribution of eight thousand diploids. To get more precise IBD segment endpoints with `tskibd`, we use 0.00001 Morgans windows, which increases runtime eightfold or more. Some true IBD segments will not be detected if the window size is too large, but decreasing the window size increases runtime.

**Table 1.** Average runtime to simulate detectable IBD segments with `ARGON` and `tskibd`

| Method | Samples | Compute Time (s) |
| --- | --- | --- |
| **ARGON**<sup>1,2</sup> | 500 | 211.80 |
| | 1000 | 652.70 ($\approx 11$ min) |
| | 2000 | 3292.90 ($\approx 55$ min) |
| **tskibd** | 500 | 6.80 |
|  | 1000 | 28.80 |
|  | 2000 | 151.78 |
| | 4000 | 921.60 ($\approx 15$ min) |
| | 8000 | 4409.40 ($\approx 73$ min) |

<sup>1</sup>Runtimes to simulate ARGs are not included in the results, which are small to negligible percentages of total runtimes.
<sup>2</sup>Sliding non-overlapping windows are of size 0.0001 Morgans.

In comparison, for eight thousand diploids, our improved approach simulates IBD segments ≥ 0.01 Morgans around a locus in less than two seconds (Figure 2). Even our naive approach completes the same scope of simulations in less than two minutes. Our method is exact for a single locus, whereas `tskibd` may be inexact due to its windowing heuristic. Moreover, our method provides the full locus-specific length distribution whereas `ARGON` and `tskibd` provide a length distribution truncated by the size of the genomic region. Conversely, our method relies on a mathematical construct without regard to finite chromosome sizes, which can result in detectable IBD lengths exceeding the chromosome size.

## 9 Discussion

To efficiently simulate IBD segment lengths overlapping a focal location, we exploit the fact that small values occur with high probability in Gamma random variables. Fast simulation in population genetics is important for statistical methods like approximate Bayesian computation (Beaumont et al., 2002), importance sampling (Browning, 2000; Stern et al., 2019), and neural network learning (Korfmann et al., 2023). Our method was developed with the evaluation of statistical consistency (Casella and Berger, 2002), parametric bootstrapping (Efron, 1987), and asymptotic distributions (Temple and Thompson, 2024) in mind.

Existing methods `ARGON` (Palamara, 2016) and `tskibd` (Guo et al., 2024) simulate IBD segment lengths for genomic region sizes less than 0.10 Morgans and thousands of samples within hours to days. These runtime performances are insufficient for the aforementioned methods and analyses, in particular parametric bootstrapping (Temple et al., 2024; Efron, 1987). We benchmark that our average runtime scales approximately linear as the number of haplotype pairs scales quadratically in sample size, taking as little as a couple of seconds or tens of seconds for sample sizes of order 10^4^ or 10^5^, respectively. The pruning and merging techniques presented here for a single locus could motivate changes to `ARGON` and `tskibd` that improve the runtime of genome-wide IBD simulations.

Related studies have already used our algorithm to these ends. Running our algorithm tens of millions of times with samples sizes ≥ 5000, Temple and Thompson (2024) show simulation results that are consistent with the conditions of their central limit theorems. Running our algorithm millions of times with a sample size of five thouand diploids, Temple et al. (2024) show that ninety-five percent parametric boot-strap intervals for a selection coefficient estimator contain the true parameter in ninety percent of simulations. They also show that exploring the effects of sample size and detection threshold on selection coefficient estimation is feasible on a laptop. Temple (2024) assesses the tradeoffs between standard normal and percentile-based confidence intervals for the Temple et al. (2024) selection coefficient estimator. Temple (2024) also shows how to calculate the statistical power in an excess IBD rate scan as the magnitude of directional selection increases. These studies would otherwise have been computationally intractable using the existing methods `ARGON` and `tskibd`. Indeed, the scope of the Temple and Thompson (2024) simulations amounts to hundreds of days of computing time even with our efficient algorithm.

The algorithm may assist in developing IBD clustering methods as well. A previously published method to simulate IBD cluster sizes comparable to those observed in human data is based solely on heuristics (Shemirani et al., 2023), whereas our method is an exact simulation of the process. Temple et al. (2024) developed their method to find abnormally large IBD clusters by experimenting with our simulations. Generating IBD clusters, which are qualitatively different from Erdos-Renyi networks (Temple and Thompson, 2024), could be fruitful in IBD network analyses (Shemirani et al., 2023). The distribution of IBD cluster sizes could also help benchmark multi-way IBD segment detection (Browning and Browning, 2024).

## Statements and Declarations

### Ethics approval and consent to participate

Not applicable

### Consent for publication

Not applicable

### Data availability

Not applicable

### Materials availability

Not applicable

### Code availability

The algorithm is in a Python package (https://github.com/sdtemple/isweep), which is available under the CC0 1.0 Universal License.

### Author contributions

S.D.T. proposed the study, planned the study, wrote the software, conducted the analysis, and wrote the manuscript. S.D.T. and S.R.B. developed the algorithm. S.D.T. and E.A.T. derived the theoretical results. All authors contributed to editing the manuscript.

## Appendix A The number of identity-by-descent comparisons

### A.1 Computations near the root of the coalescent tree

#### Lemma 3.

*Recall that B*_*j*_ ∼ *Binomial*(*B*_*j*−1_, 1*/*2) *for j* ≥ 1 *and B*_0_ = *n. Then*,

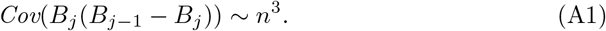

*Proof*. We apply the laws of total covariance and expectation in a recursive fashion. Overall, we must control three terms:

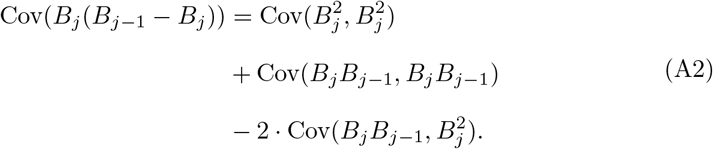

The first four moments of the conditional binomial random variable are useful:

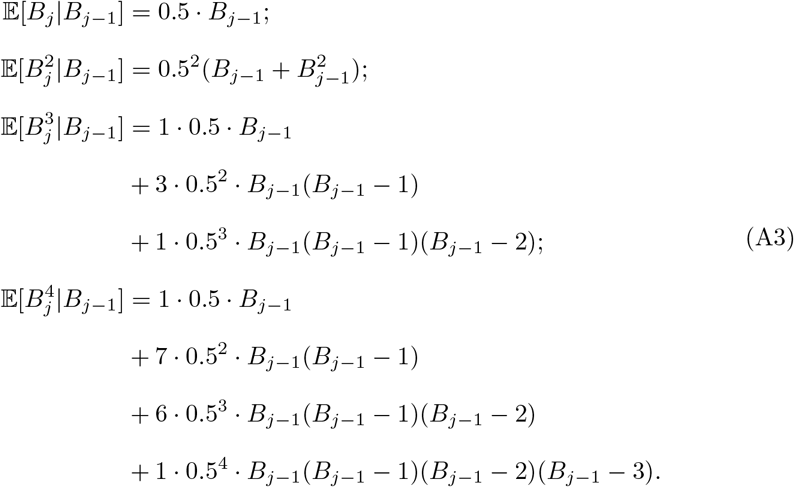

The following conditional covariances are also useful.

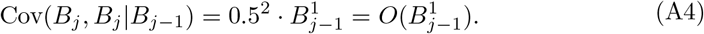

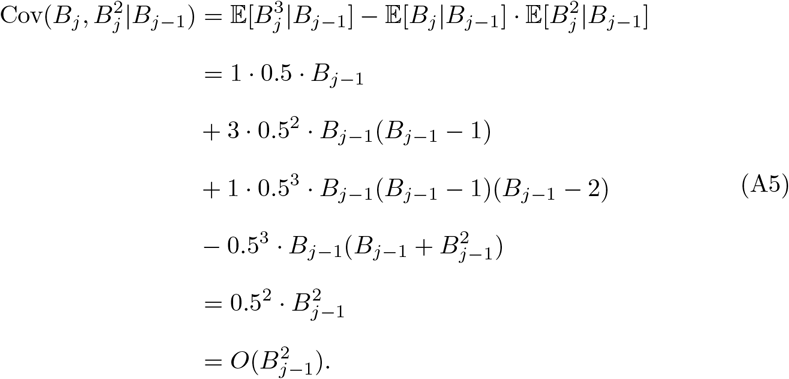

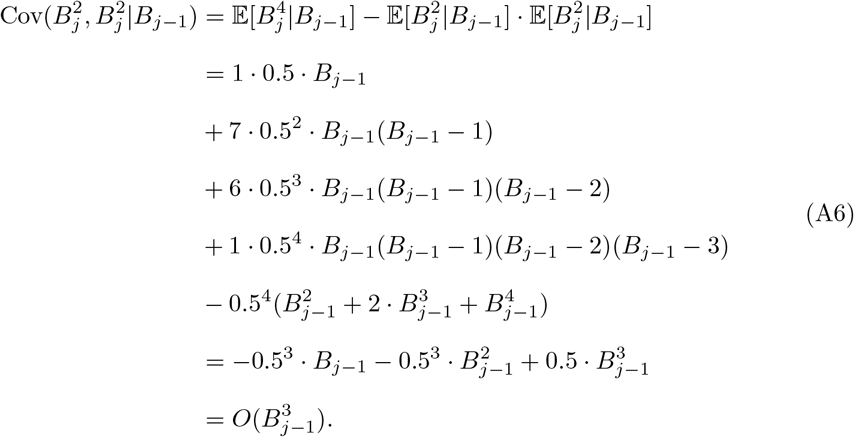

Notice that all of the conditional covariances are of an order of three or less. By recursively applying the law of total expectation, we derive 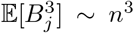.Another important unconditional covariance term is

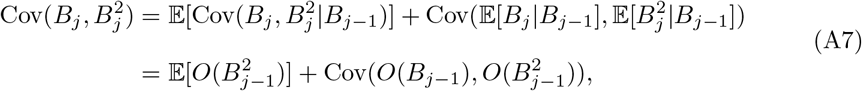

which is asymptotically equivalent to *n*^2^ when the total laws of expectation and covariance are applied recursively. Finally, we evaluate the asymptotic behavior of the three unconditional covariances in Equation A2.

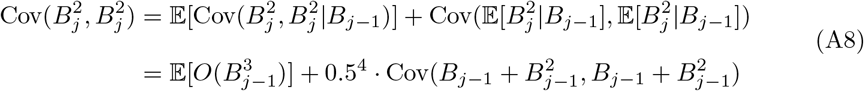

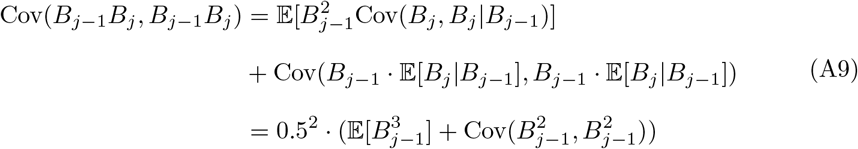

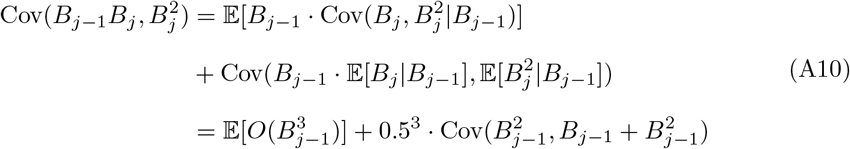

By recursively applying the total laws of expectation and covariance, we conclude that Equations A8, A9, and A10 are asymptotically equivalent to *n*^3^.

### A.2 Computations near the leaves of the coalescent tree

A complementary perspective on joint subtree sizes we take from Dahmer and Kersting (2015). Now, we work upward from the leaves to the root. Dahmer and Kersting (2015) provide a lemma for the expected number of subtrees containing *r* sample haplotypes at the (*n* − *k*)^th^ coalescent event. We can use their moment calculations together with the expected value of the hypoexponential random variable 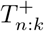 to build intuition for the average subtree sizes at a specified generation. In a toy example, Figure S7 shows the average number of sample haplotypes under subtrees of a given size at generations 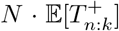.Our main observation is that before the final coalescent events, which occur at expected times proportional to population size *N*, most sample haplotypes are expected to be under a subtree of size an order of magnitude smaller than the sample size.

## Supplementary figures

**Fig S1.**
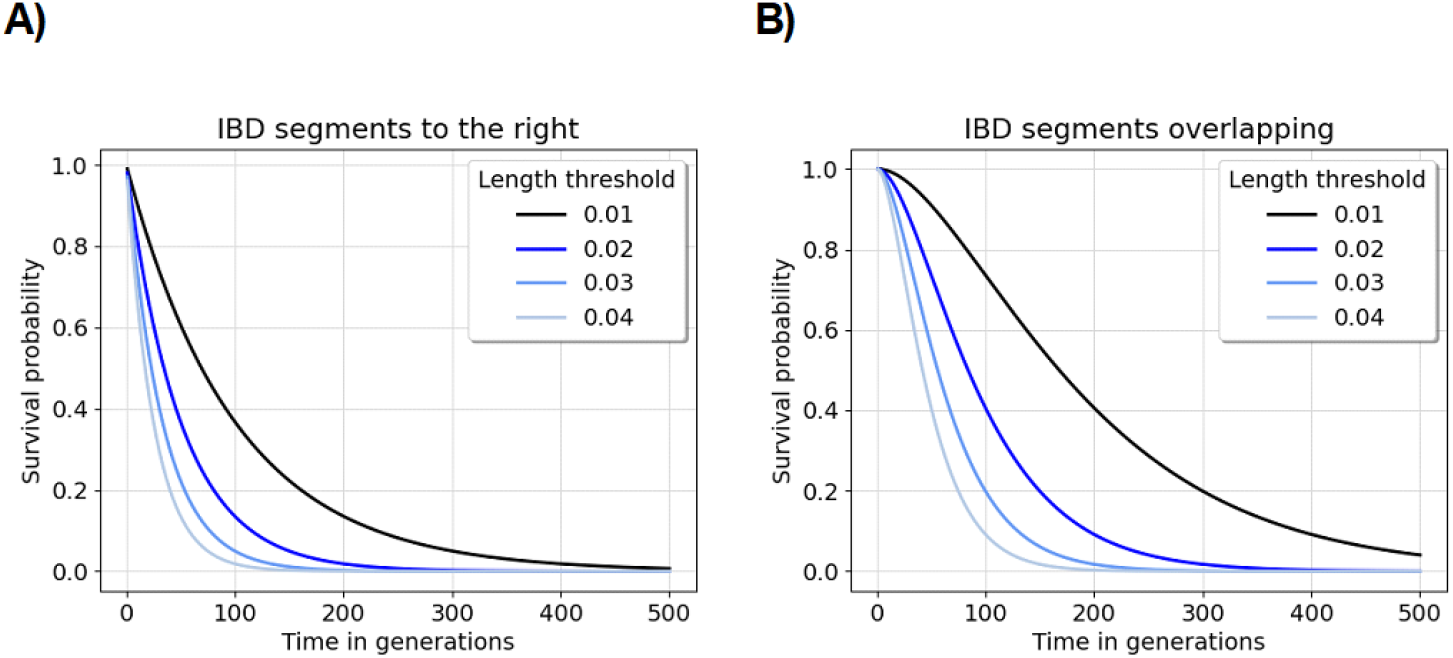
The upper tail probabilities of Gamma random variables. Subplots A) and B) show the survival probabilities for shape parameters 1 and 2, respectively. The rate of the random variables is the coalescent time in generations (*x*-axis). The survival probability (*y*-axis) comes from Equations 6 and 7. The length thresholds are denoted by different colors and line styles, defined in the legend.

**Fig S2.**
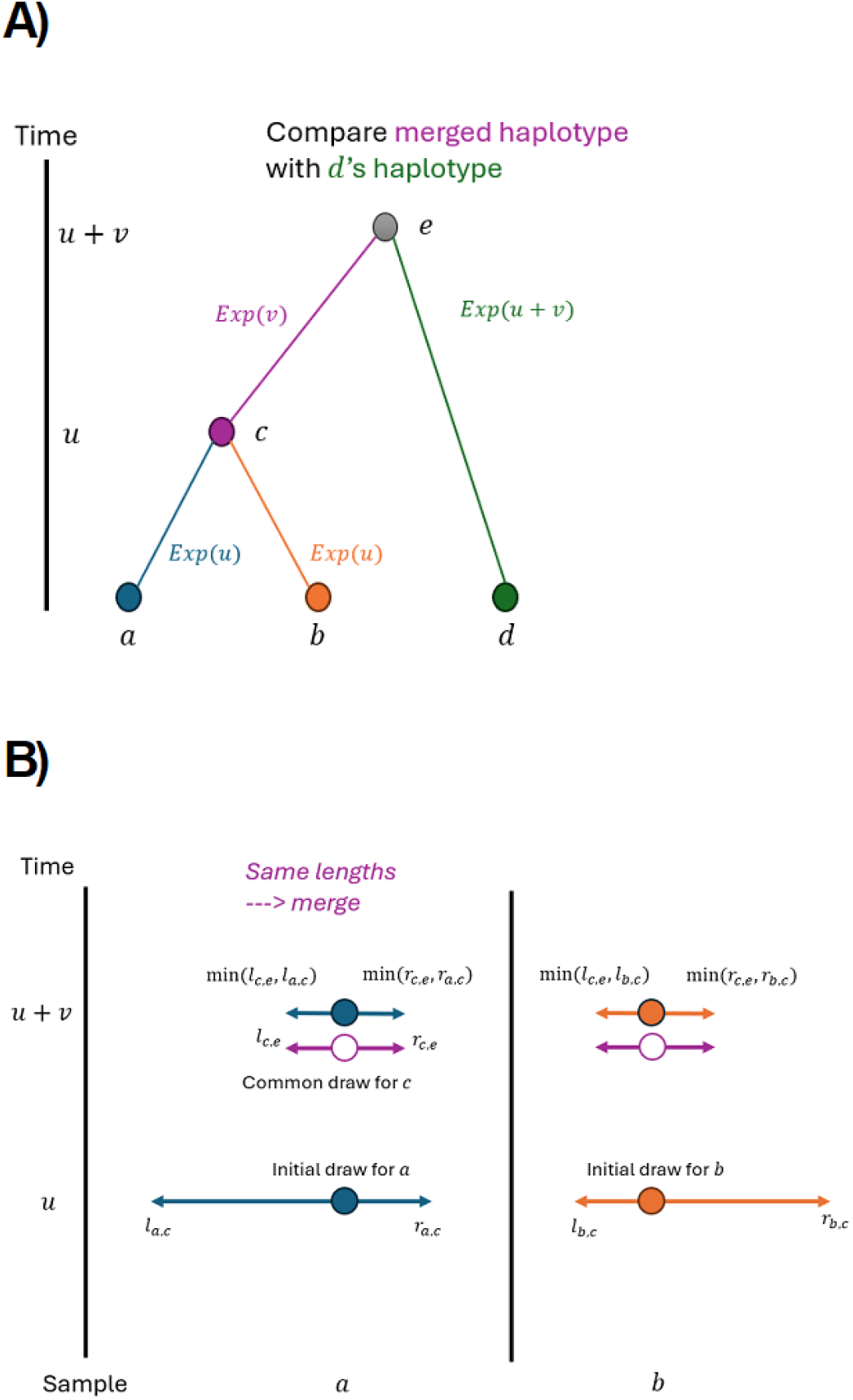
Illustration of merging haplotypes. A) We draw recombination endpoints to the left and right of the focal location from Exponential(*u*) for both sample haplotypes *a* and *b* at coalescent time *u*. We draw recombination endpoints to the left and right of the focal location from Exponential(*v*) for the common ancestor *c* of *a* and *b* at coalescent time *u* + *v*. Colors denote branch lengths and recombination endpoints corresponding to a given haplotype. *l*_*a,c*_ and *r*_*a,c*_ denote the endpoints for *a* drawn to the left and right of the focal location at time *u* (lowercase denotes observation of random variables). B) We compute minimums of lengths drawn for *c* and *a* and *c* and *b*, respectively. We merge the sample haplotypes *a* and *b* in future calculations because the minimum lengths are both the recombination endpoints drawn from the common ancestor *c*. When comparing recombination endpoints at time *u* + *v* with those of the haplotype *d*, we make one comparisons. The haplotypes remain longer than the detection threshold *w* Morgans.

**Fig S3.**
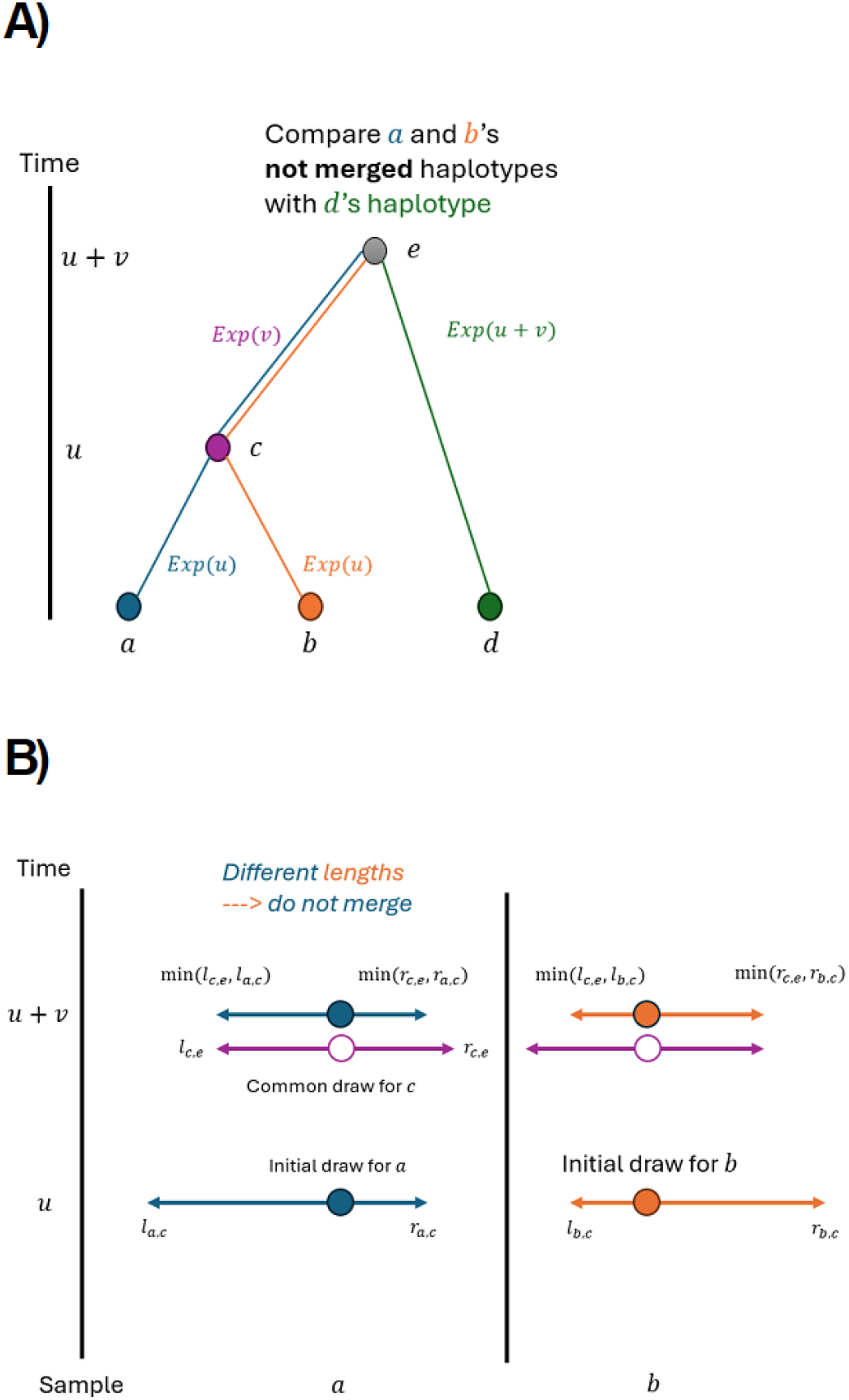
Illustration of ***not*** merging haplotypes. A) We draw recombination endpoints to the left and right of the focal location from Exponential(*u*) for both sample haplotypes *a* and *b* at coalescent time *u*. We draw recombination endpoints to the left and right of the focal location from Exponential(*v*) for the common ancestor *c* of *a* and *b* at coalescent time *u* + *v*. Colors denote branch lengths and recombination endpoints corresponding to a given haplotype. *l*_*a,c*_ and *r*_*a,c*_ denote the endpoints for *a* drawn to the left and right of the focal location at time *u* (lowercase denotes observation of random variables). B) We compute minimums of lengths drawn for *c* and *a* and *c* and *b*, respectively. We ***do not*** merge the sample haplotypes *a* and *b* in future calculations because the minimum lengths are ***not*** both the recombination endpoints drawn from the common ancestor *c*. When comparing recombination endpoints at time *u* + *v* with those of the haplotype *d*, we make ***two*** comparisons. The haplotypes remain longer than the detection threshold *w* Morgans.

**Fig S4.**
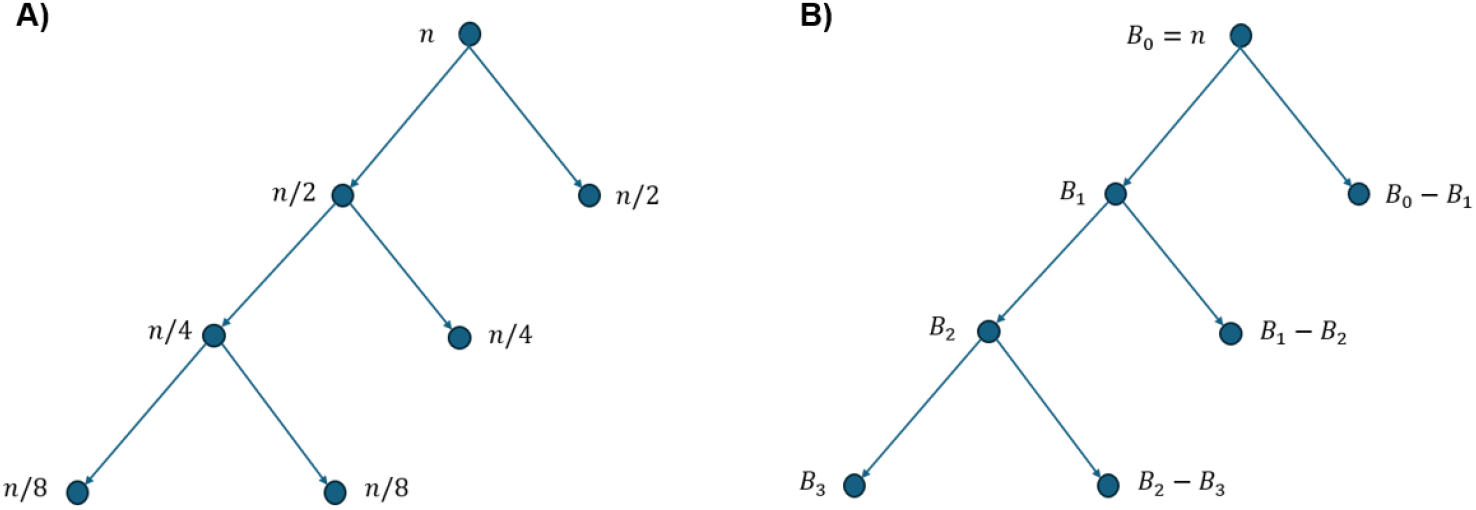
Illustration of worst-case subtree sizes in random bifurcating tree. A) The worst-case subtree sizes of ancestors (dots) are when each bifurcation is an even split. B) The model {*B*_*j*_} of subtree sizes down one branching path is defined as random variables. The sample size is denoted as *n*.

**Fig S5.**
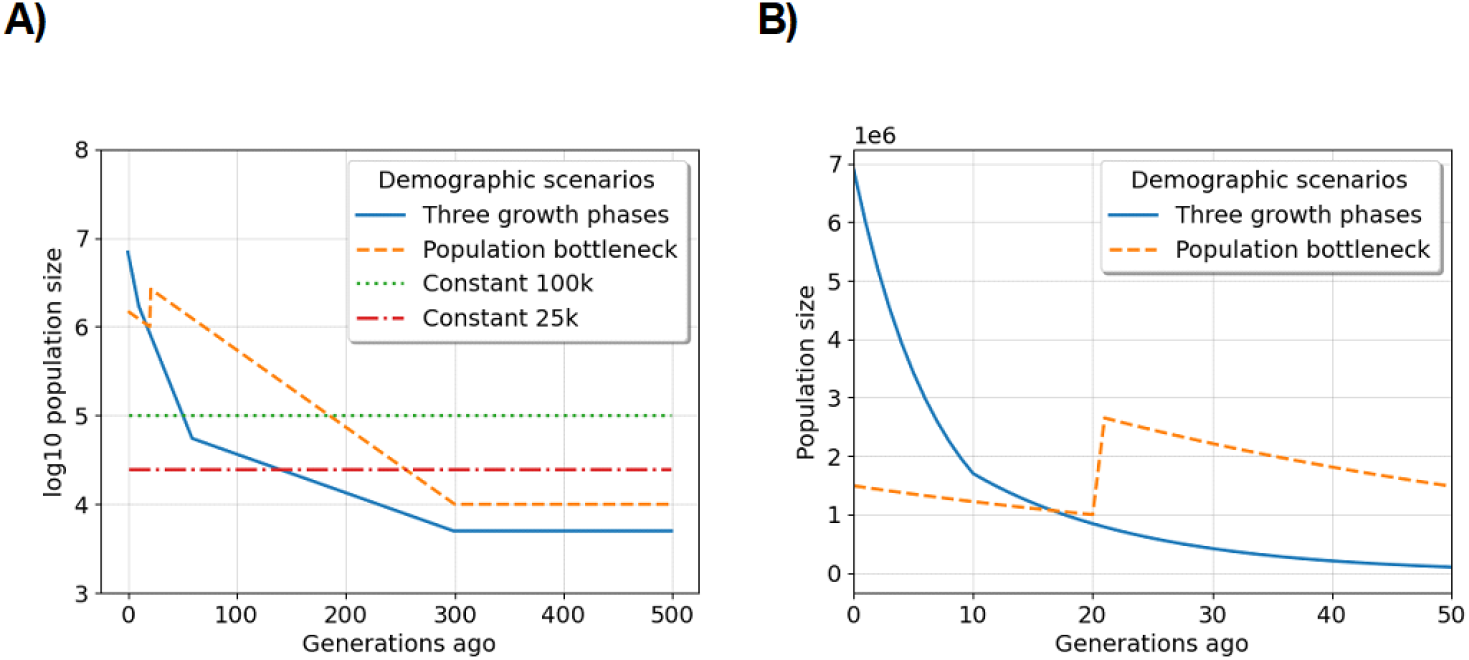
Demographic scenarios we consider in simulation studies: A) coalescent time in generations ago by the log 10 population size, and B) the most recent fifty generations by population size for examples of exponential growth. The legends specify the color and line style for each scenario. As opposed to coalescent time used in the main text, we describe the scenarios moving forward in time here. The three phases of exponential growth model is as follows: a population of ancestral size five thousand diploids increases exponentially each generation at rates one, seven, and fifteen percent starting three hundred, sixty, and ten generations ago. This demographic model is similar to the “UK-like” model in Cai et al. (2023). The population bottleneck model is as follows: a population of ancestral size ten thousand diploids increases exponentially each generation at a rate of two percent starting three hundred generations ago, but twenty generations before the present day, the population experiences an instantaneous reduction in size to one million diploids. Otherwise, the demographic scenarios we explore here are populations of constant size twenty-five and one hundred thousand diploids.

**Fig S6.**
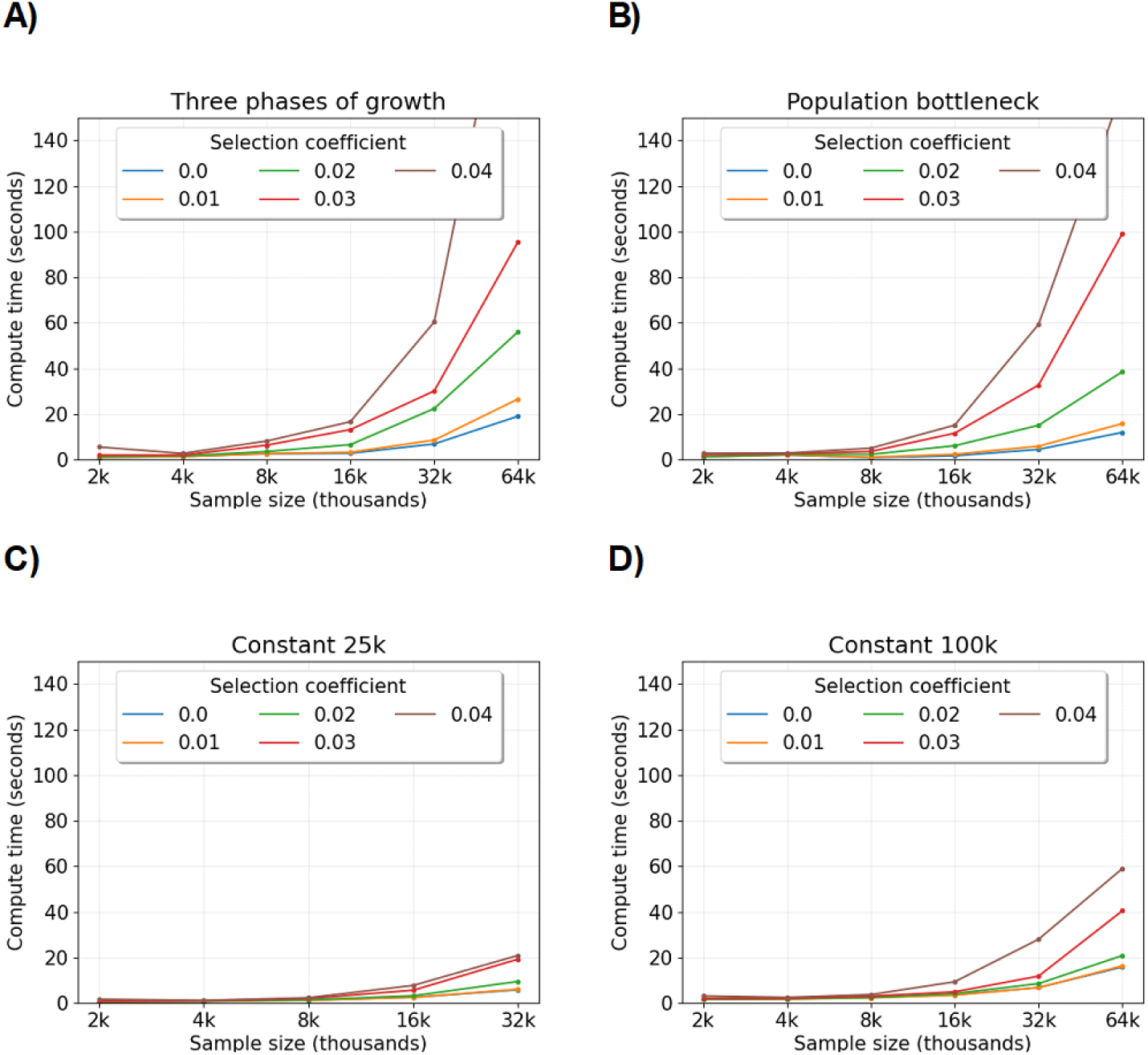
Compute time to simulate IBD segment lengths around a locus depending on demography and selection. Compute time (*y*-axis) in seconds by sample size (*x*-axis) in thousands is averaged over five simulations. The legends denote colored line styles for different selection coefficients. A), B), C), and D) show results for demographic scenarios of three phases of exponential growth, a population bottleneck, and constant population sizes of twenty-five and one hundred thousand diploids, respectively. The Morgans length threshold is 0.01.

**Fig S7.**
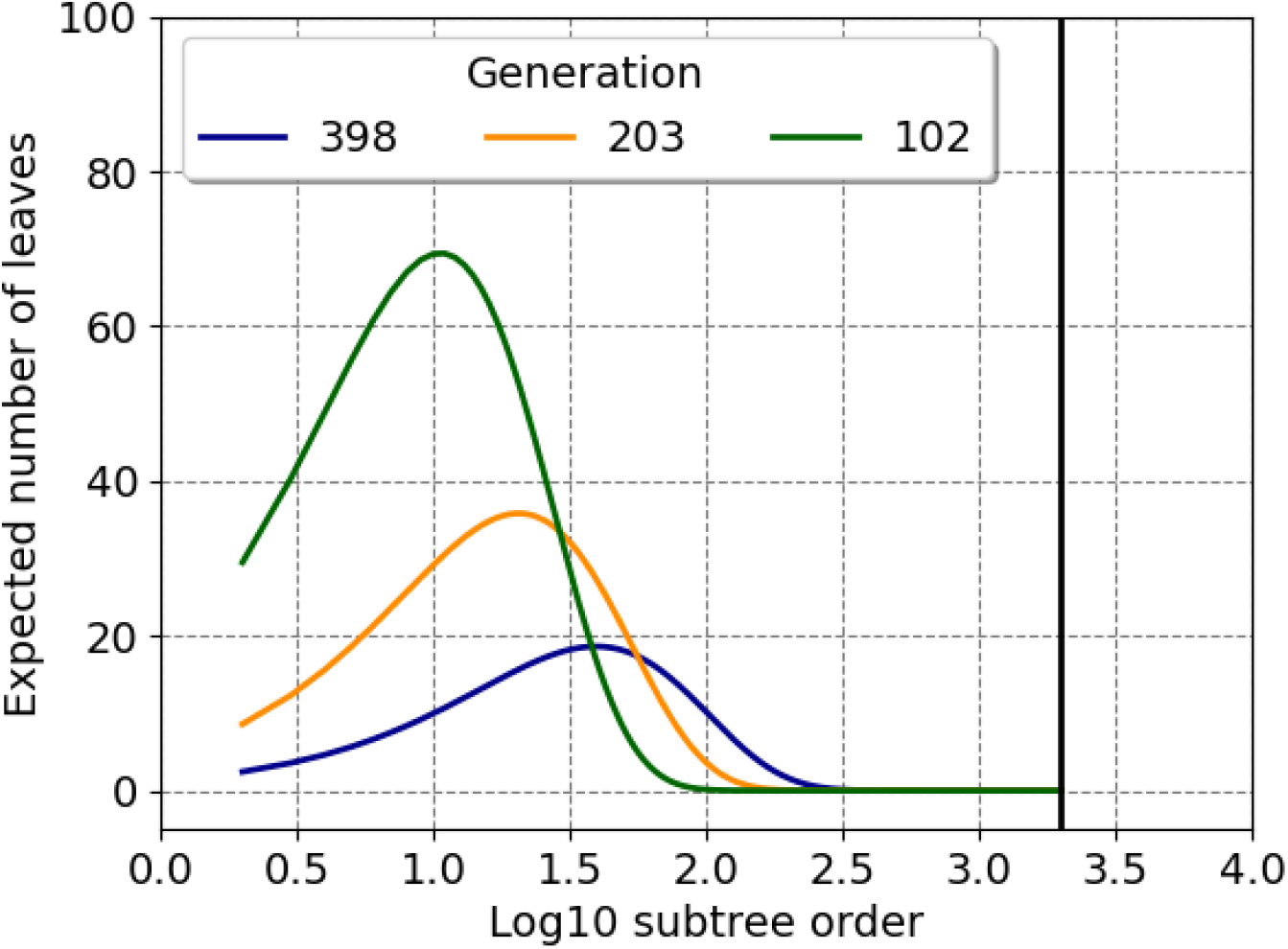
The expected cardinality of subtree sizes at different coalescent times. Using Lemma 1 in Dahmer and Kersting (2015), we compute the expected number of subtrees containing *r* samples (*x*-axis) at the (*n™ k*)^th^ coalescent event. We multiply these moments by *r* to get the expected number of leaves under such subtrees (*y*-axis). There are two thousand samples. Dark blue, orange, and green lines correspond to *k* =50, 95, and 180. We compute the expected time of the (*n™ k*)^th^ coalescent event (Hein et al., 2005) and multiply by a population size of ten thousand to get generations (legend). The vertical line is logarithm 10 of sample size.

